# Collagen-induced inflammations

**DOI:** 10.1101/2020.11.09.373761

**Authors:** Ningchao Du, Yang Li, Min Xu, Changlin Wu, Xiaolan Chen, Feng Yang, Qiong He, Muchun Liao, Jianfu Qiu, Changhao Wang, Jun Sun, Xiang Ma, Guang Sheng, Kun Du, Kutty Selva Nandakumar, Li Song

**Author notes:** Corresponding Author: Li Song.

## Abstract

Collagen-induced arthritis (CIA) mouse model is currently the most widely used and reliable autoimmune model to study rheumatoid arthritis. In this model, we used bovine type II collagen (CII) and complete Freund’s adjuvant (CFA) or incomplete Freund’s adjuvant (IFA) to form emulsifier, and mice were injected intradermal to induce autoimmune arthritis (CIA). In this model, we ground bovine collagen type II (CII) with complete Freund’s adjuvant (CFA) or incomplete Freund’s adjuvant (IFA) to form emulsifiers, and intradermal injection in mice induced autoimmune arthritis (CIA). This article describes the mouse, CFA strains, key emulsification, anesthesia, and injection immune techniques, as well as the incidence, date of onset, score, pathological results of arthritis. The total time for preparation of reagents and immunization of 20 mice was about 2-2.5 hours. In this protocol, we induced a high incidence of CIA with DBA/1J in genetically susceptible mice and assessed the severity and pathology of the disease, at the same time we found that CII also can induced enteritis, including ileitis and colitis. The initial symptoms of arthritis appear in the 24-26 days of the experiment, that is, 3-5 days after the second immunization, the peak period of inflammation was 30-36 days, the arthritis incidence about 90-100%, at the same time, the incidence of enteritis and arthritis were the same, small intestinal inflammation was more severe, but the duration was short; while the colonic inflammation was mild, and the duration was longer than enteritis, we named it collagen induced inflammations (CIIs).

## 1. INTRODUCTION

Rheumatoid arthritis (RA) is a chronic autoimmune inflammatory disease, which initially manifests as joint swelling, pain, stiffness, etc., and eventually leads to the erosion of cartilage and bone tissue through long-term invasion of vascular fibers [1]. RA is characterized by persistent synovitis, systemic inflammation, and autoantibodies. The pathological process of RA is long and complex, including synovial cell proliferation, pannus formation, cartilage and bone erosion. Rheumatoid arthritis accounts for 0.5-1.0% of adults, with 5-50 cases per 100,000 new cases per year, the quality of life and life of patients are seriously threatened, so it is worth our in-depth study. Collagen-induced arthritis (CIA) mouse model is currently the most widely used and reliable autoimmune model to study RA. Similarities between CIA and RA are both autoantibodies that produce collagen and induce inflammation through autoimmune responses, and collagen is also one of the important autoantigens observed in human RA [2–4]. Understanding the process of pathogenic RA In humans, rodent models are classical animal models. These animal models can test the efficacy, efficiency and safety of drugs, etc. It is of major significance for the development of new drugs and the discovery of new targets.

CIA has been studied more extensively as animal model because it has many similar pathologies and immunological characteristics to human RA [5]. In CIA model, an immune response is being directed against a joint antigen such as collagen type II (CII) [6]. CII comes from a variety of sources such as bovine, humans, pigs, and chickens, and responses are thought to be different, it also needs its own T, B cells and cytokines to participate in the immune response to CII. First CIA model was established by immunizing rats at CII [7]. Later, CIA models were replicated mice and monkeys respectively. There are some mice strains that do not respond well to CII. The susceptible strains including DBA/1, B10.Q and B10.RIII etc, DBA/1 (H-2q) mice is used widely as the CIA model now [8], are highly susceptible to CIA and respond to chick, bovine, and porcine CII.

At present, there are few reports about other inflammation induced by CII, such as enteritis, etc. We found that CII can also cause intestinal inflammation, while the severity and duration of inflammation in the small and large intestine are different, which needs further study.

## 2. MATERIALS AND PROCEDURE

### 2.1 MATERIALS

#### 2.1.1 REAGENTS

. Immunization Grade Bovine Type II Collagen (CII, Chondrex, cat. no.20021, 10 mg/ea, lyophilized).
. Complete Freund adjuvant (CFA, Sigma, cat. no. F5881-10ML, 10 ml/ea), including heat-killed mycobacterium tuberculosis strain H37Ra (ATCC 25177).
. Incomplete Freund adjuvant (IFA, Sigma, cat. no. F5506-10ML, 10 ml/ea).
. Isoflurane (BD, R510).
. Experiments DBA/1J mice conformed to national and local regulations.

#### 2.1.2 EQUIPMENT

. Mortar and pestle (A ceramic mortar with an inner diameter of 50 mm is enough to emulsify CII for 20 mice at one time.)
. 1ml syringe (Kangli, #KL001)
. 50ml test tube (Corning, #358206)
. Appropriate amount of test ice

### 2.2 PROCEDURE

#### 2.2.1 Preparation and storage of CII

Solid bovine collagen type II (CII, Chondrex, 20021) keep in −20 °C refrigerator, was dissolved in 0.01M acetic acid, shake for 3-5 minutes at room temperature to dissolve CII sufficiently to prevent CII emulsifier concentration error caused by incomplete dissolution of CII. Store it in 4 degree refrigerator and use it separately.

CAUTION Do not prepare too much CII mother liquor at one time to prevent CII deterioration.

#### 2.2.2 Emulsification of CII. TIMING 60-90 min

Bovine collagen type II (CII, Chondrex, 20021) took from −20°C and was dissolved in 0.01M acetic acid keep it in 4°C.When use it and added CII to CFA/IFA in mortar dropwise on ice, mortar should be kept cool on ice 30 minutes ago to avoid damage to CII at room temperature, emulsified in equal volume with CFA containing 1mg/ml heat-inactivated mycobacterium tuberculosis (H37Ra, ATCC 25177), or emulsified with IFA in equal volume, keeping their ratios at 1:1 (Fig.1A).

**Fig. 1.**
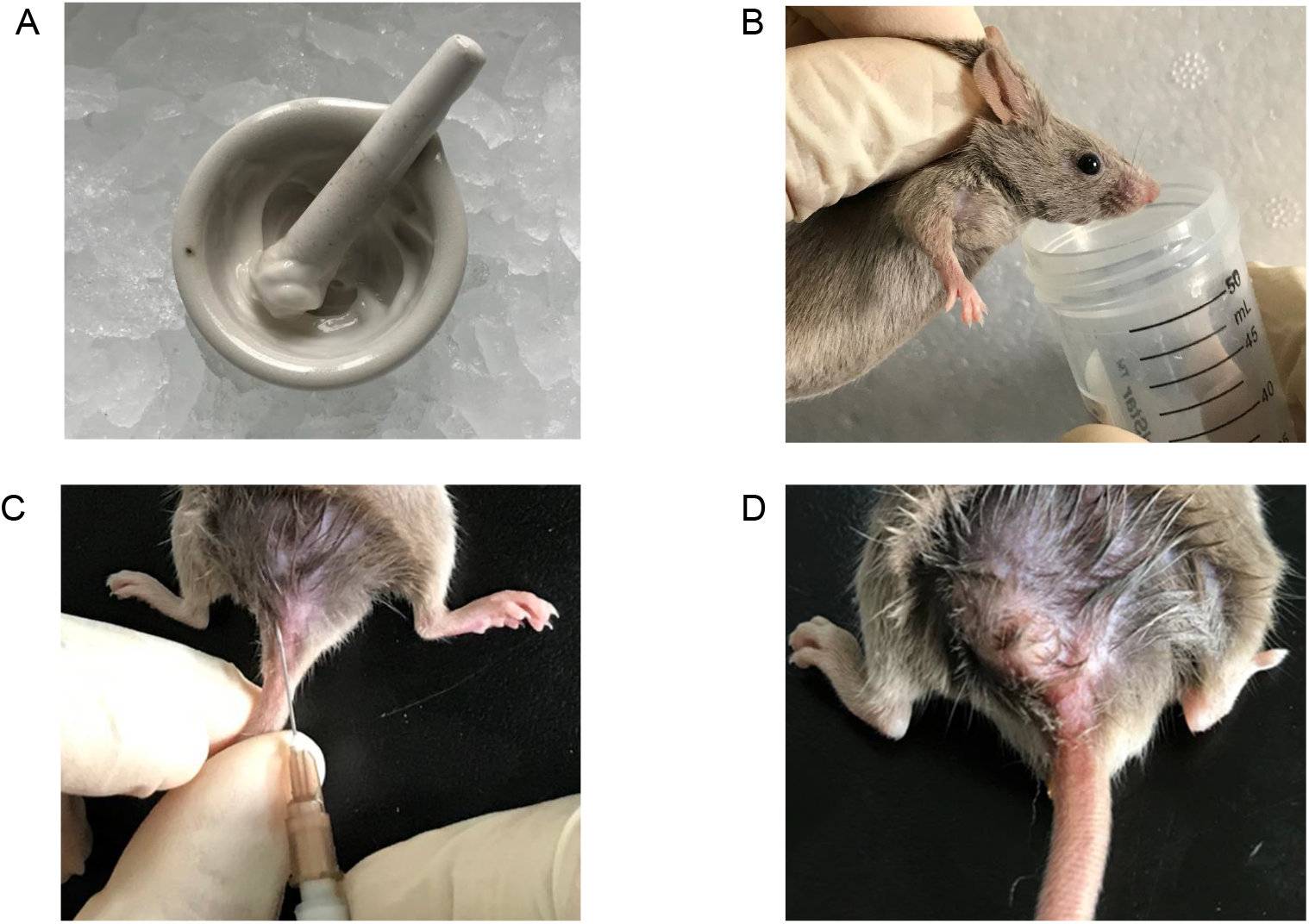
CII Emulsification, anesthesia and injection immunization for Mice. (A) Emulsification process of CII and adjuvant. (B) The process of anesthesia in mice. (C) Tail root injection in mice. (D) Pictures of mice after subcutaneous injection of tail roots.

Emulsification should pay attention to the indoor temperature and humidity. If the temperature is too high, the emulsion is easy to be stratified; the humidity should be kept at 50-60%. If the humidity is too high, it will easily cause the mortar inner wall to liquefy and form water droplets, which will cause the emulsion to be diluted, thus leading to immune failure. When the emulsion droplets do not disperse on the water surface, it means emulsify completely and they can be injected with immunization. The emulsion should be put on ice and used up within 60 mins. If the time is too long, the CII will decompose or the liquid will be stratified, leading to immune failure. Emulsifiers drawn in 1 ml syringe should also be placed on ice. The amount of emulsification prepared each time should be 0.4-0.5 mL more than the calculated amount, that is, the amount of 4-5 more mice, in addition to the loss is basically enough.

CAUTION During emulsification, CII was added drop by drop to the adjuvant surface which was put into the pre-cooled mortar in advance, stirred to make it fully mixed, and grinded on ice as soon as possible. Grinding in a clean environment should not be affected by environmental factors leading to other immune reactions.

#### 2.2.3 Anaesthesia of mice. TIMING 20-30s

Mice can be anesthetized by inhalation for 3-5 seconds after obvious muscle relaxation. The total anesthesia time is about 20-30 seconds. Extended anesthesia time is prone to death, and even chest compression can not save their lives (Fig.1B).

CAUTION When anesthetizing mice, attention should be paid to the grip mode of mice, so as not to make their neck strangled and lead to hypoxia, otherwise it is more likely to cause anesthesia lethality.

#### 2.2.4 Preparation of CII. TIMING 1-2 min

Inhale CII emulsion into a 1 ml syringe, force the emulsion to the end of the syringe, and push the syringe piston forward to expel air, then put on ice prepare to use.

#### 2.2.5 Immunization of mice. TIMING 1 min per mouse

For the first immunization of 0 day this emulsion were injected i.d. part of tail root with 100ul / CII 100ug / mice. The booster injection were given on 21 days, CII was emulsified in equal amounts with IFA, this emulsion were injected the other i.d. side of the tail root with 100ml / CII 50ug / mice [9]. The tail roots of mice were disinfected with 75% alcohol and injected intradermally (i.d.) into one side of the tail roots (Fig. 1C), 0.1 ml/mouse, form a white mass (Fig. 1D).The white mass can be seen after intradermal injection of CII emulsion. If there are only bumps on the surface of the skin after injection but not white, the injection may be too deep and enter the subcutaneous. The other side of the tail root skin was reserved for the second immunization. After 1-2 weeks, the subcutaneous tissues will wrap up and form a mass basically leading to immune failure. Even the bump was gently crushed and can not play the immune response of slowly releasing CII.

CAUTION In many cases, CFA can not be used for the second immunization, because it will lead to excessive immune response and cause ethical and mouse death and other issues. Pay attention to dissolving CII with 0.01M glacial acetic acid, if the pH value of the solution is too low, emulsification will fail, or the mice will die soon after injection.

#### 2.2.6 Evaluation arthritis score and incidence. TIMING 5–8 weeks

To observe the skin condition at the injection site of mice after immunization, if there is a small ulcer, sterilize the injection site with 75% alcohol cotton ball, twice a day, about 1 week can basically heal. If the ulcer continues to enlarge or form infection foci, the mice should be excluded from the group and killed.

The arthritis is markedly red and swollen, with limited activity, and identification is not difficult. In the later stage of recovery from arthritis, the redness and swollen subside. Sometimes identification is difficult. As described by Nanakumar etc before [10], scoring was done blindly by a scoring system based on inflamed joints in each paw, inflammation was defined by swelling and redness of jionts. In this scoring system, each inflamed toe or finger joint has a maximum of 1 point, 5 points for each paw, 5 points for metacarpal or metatarsal bones, and 5 points for wrists and ankles. Thus, the maximum score per paw was 15, and the maximum score per mouse was 60. We scored the arthritic paws of each mouse and recorded them every 3 days.

#### 2.2.7 Histopathology and micro-CT analysis

The paws of mice were fixed with 4% paraformaldehyde (pH 7.0) for 24 h, decalcified for 4 weeks, and embedded in dehydrated paraffin. Sections of 6 μm were stained using hematoxylin-eosin to examine cellular infiltration and bone/cartilage morphology. Histological scores were calculated blindly using the method described earlier [11]. On the 81st day of the experiment, the bone parameters of the posterior limbs were analyzed by micro-CT. The paws of four groups of mice were scanned with Siemens Inveon Hybrid Micro-CT Scanner. Reconstruction of trabecular bone mineral content (BMC). The intestinal canal were fixed with 4% paraformaldehyde (pH 7.0) for 24 h, embedded in dehydrated paraffin, others steps same with paws.

### 2.3 Statistical analysis

Statistical analyses were performed using SPSS software version 21.0 (SPSS Inc, Chicago, IL). One-way analysis of variance (ANOVA) following LSD (L) was used for multiple range tests and two-tailed Student’s t tests were used for two group comparisons. All data were presented as mean ± SD, p < 0.05 was considered statistically significant (*, p < 0.05; **, p < 0.01; ***, p < 0.001).

## 3. RESULTS

### 3.1 The arthritis incidence and score

CIA1 and CIA2 groups have the serious arthritis compared with Con group, we can got the conclusion that all the paws maybe have the arthritis, including the forepaws (Fig. 2A). Arthritis of CIA1 and CIA2 occurred on the 25th and 26th days of the experiment, respectively, that is, the 4th and 5th days after secondary immunization.The incidence of arthritis in both groups was 100% and 90% after 31 and 30 days, respectively(Fig. 2B). Arthritis began to appear on the 24th day of the experiment, and the peak appeared on the 33rd day (Fig. 2C). There was no significant difference between CIA1 and CIA2 groups.

**Fig. 2.**
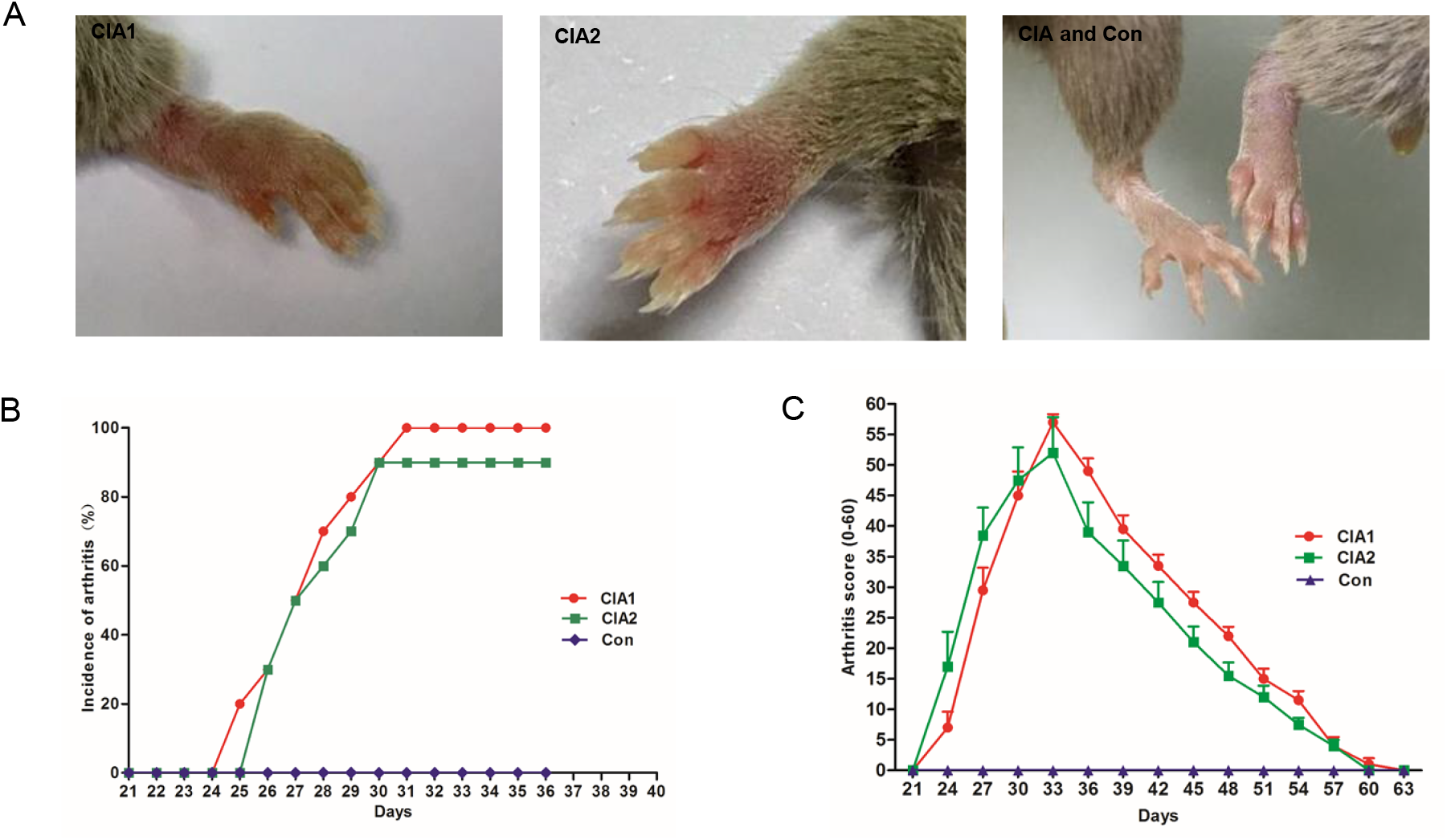
Incidence and scoring of arthritis at the peak of arthritis in CIA model. (A) Arthritis pictures from different groups of mice. (B) Onset day of arthritis (n = 10). (C) Arthritis score after first immunization (n = 10). CIA: collagen induced arthritis, Con: Control group. Data are presented as the mean ± SD in (B, C).

### 3.2 The HE staining at peak and convalescence of arthiritis

The HE staining pictures of hind paw of mice at the peak of arthritis, there were many inflammatory cell infiltration in the joint cavity of CIA1 and CIA2 groups (Fig. 3A). The HE staining pictures of hind paw of mice at the convalescence of arthritis, In CIA1 and CIA2 groups, arthritis cells decreased, but bone and cartilage were destroyed (Fig. 3B). Compared with Con group, the HE score of CIA1 and CIA2 was significantly different, but there was no significant difference between the two groups (Fig. 3C-D).

**Fig. 3.**
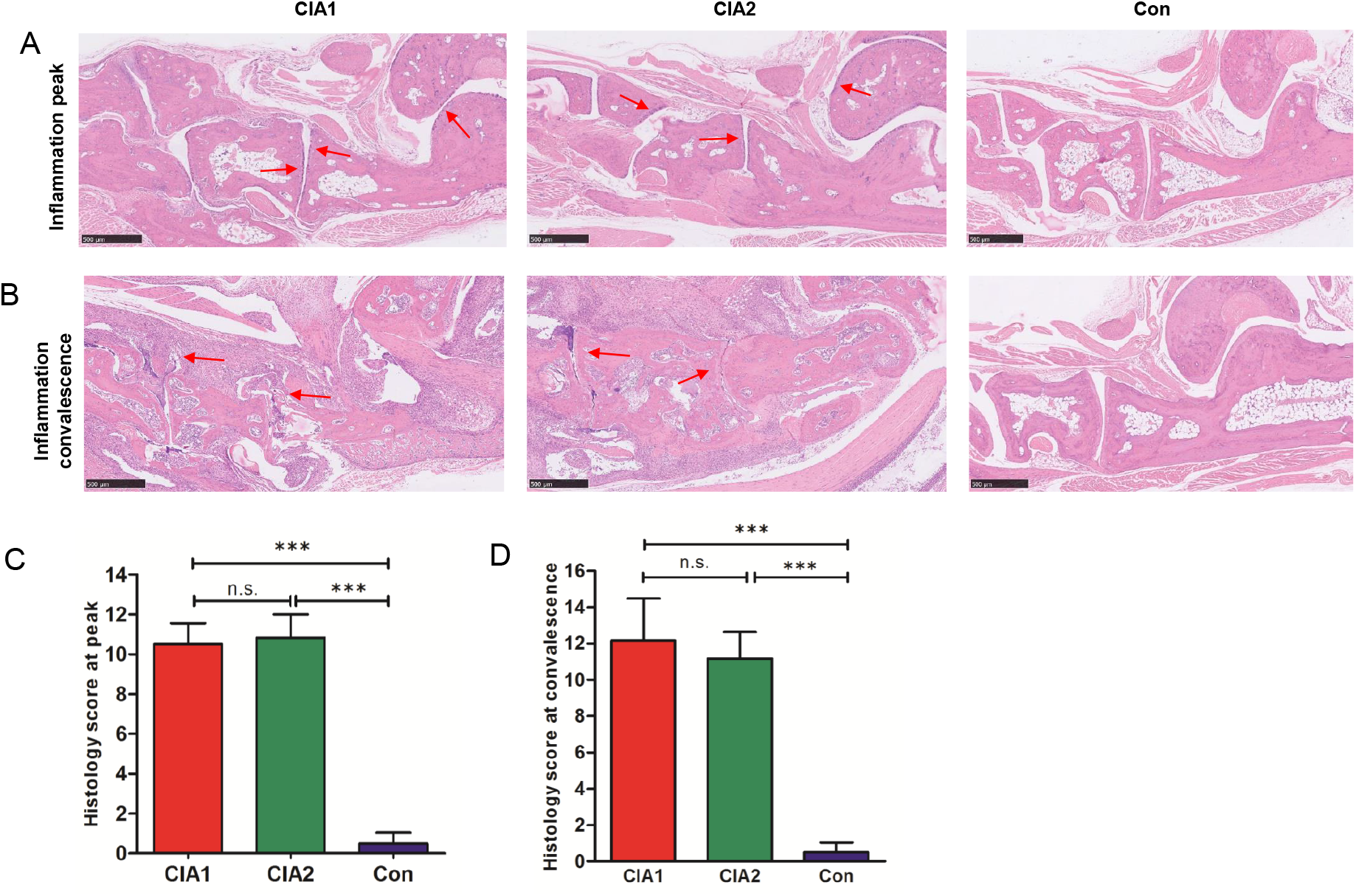
HE staining in the peak and convalescence stages of arthritis. (A) HE staining at the peak of arthritis (n = 6). (B) HE staining at the convalescence of arthritis (n = 6). (C-D) The statistical difference of HE staining at different time in CIA model. Scale bars, 500 μm. Data are presented as the mean ± SD. *** p < 0.001, n.s. stand for no significance; one way ANOVA following LSD (L) multiple range test was used in the figures (C and D).

### 3.3 The bone and joints distruction at convalescence of arthiritis

We used micro-CT scans of the hind paws of convalescent mice to investigate bone and joints distruction. The results showed that the cortical bone and joint injuries in the paws of the CIA1 and CIA2 groups were more severe than Con group (Fig. 4A). Bone morphometric parameters of BMC in CIA1 and CIA2 groups were lower than Con group, but there was no significant difference between CIA1 and CIA2 group (Fig. 4B).

**Fig. 4.**
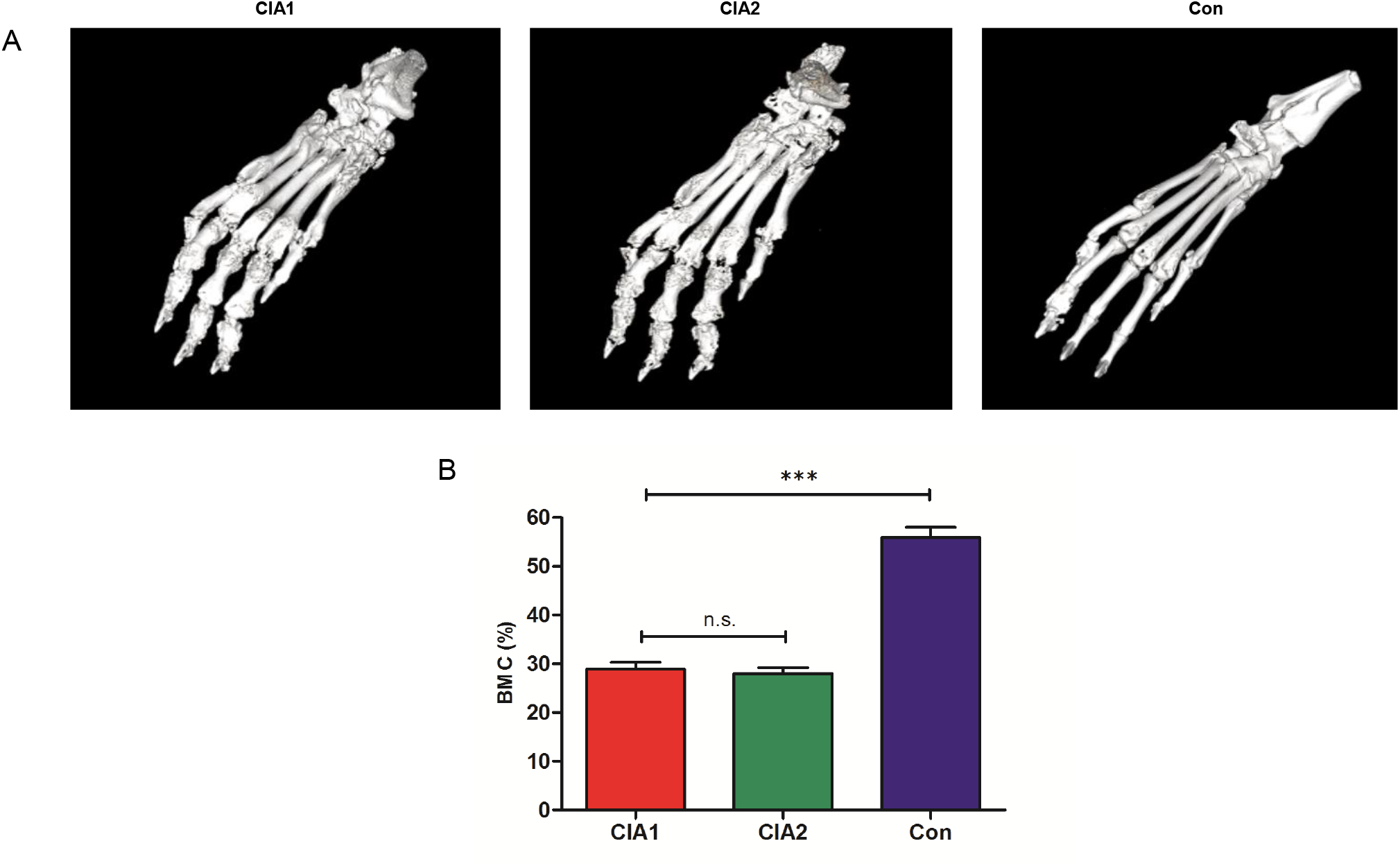
Bone and joints of different group mice at the convalescence phase of arthritis. (A) Micro-CT scanning of paws from different groups, n = 5. (B) Bone reconstruction parameters of paws from different groups. Bone mineral content (BMC) was shown. *** p < 0.001, n.s. stand for no significance; One way ANOVA following LSD (L) multiple range test (B) was used in the figure.

### 3.4 The enteritis between CIA and Con group

At the peak of CIA inflammation, all mice with arthritis had enteritis and colitis, with relatively severe enteritis and mild colitis (Fig. 5A). During the recovery period, intestinal inflammation recovered completely, while 77-80% mice still had colitis (Fig. 5B).

**Fig. 5.**
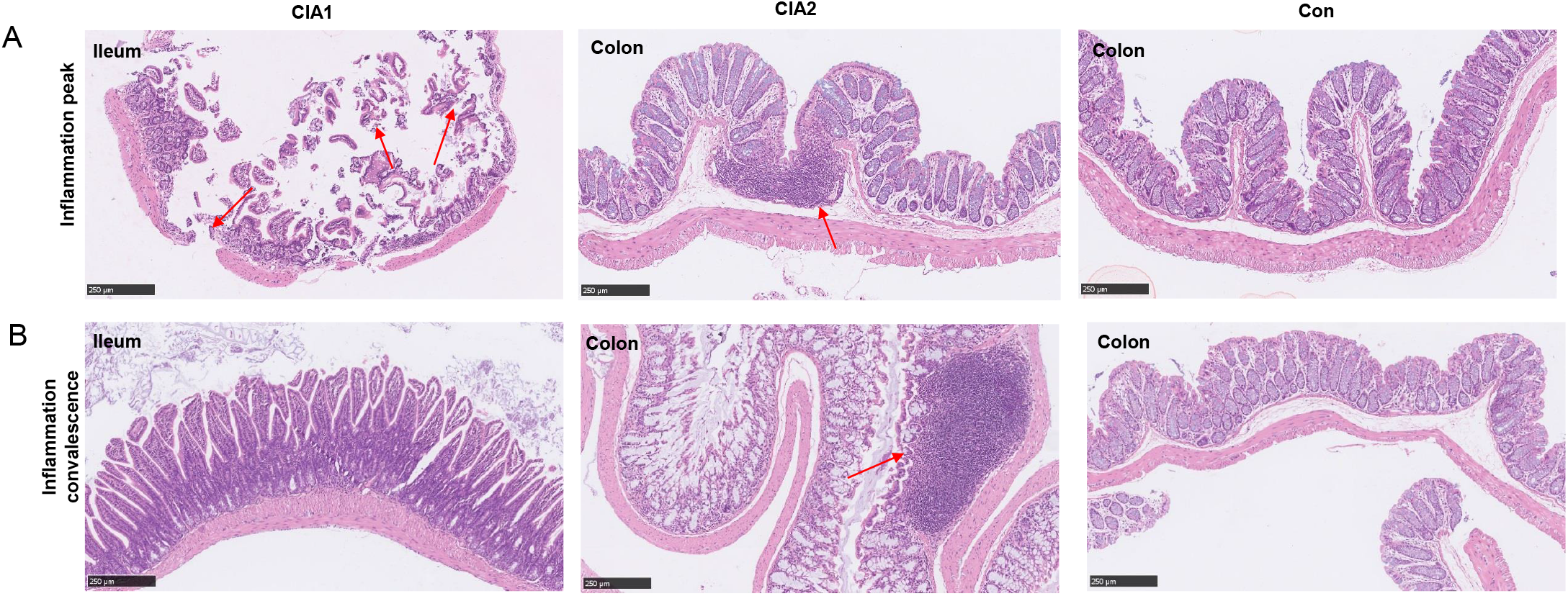
The difference of enteritis in CIA model. (A) HE staining of enteritis at peak of inflammation. (B) HE staining of enteritis at convalescence of inflammation. Scale bars 500 μm, n = 6. Data are presented as the mean ± SD.

## 4. DISCUSSION

In CIA model, after emulsification with adjuvants, slow release of CII can induce autoimmune arthritis, leading to bone and cartilage destruction and ultimately joint destruction, its pathogenesis has also been studied more clearly, while intestinal inflammation and its flora changes have been rarely studied. Microbiota has been observed has relation with early rheumatoid arthritis, the intestinal microbiota of rheumatoid arthritis was characterized by an expansion or decrease of gut microbiota. Maeda Y etc [12] have demonstrated that the abundance of *Prevolella copri* was increased in some early RA, *Prevolella histicola* from human gut microbiota suppressed the development of arthritis. In summary, *Prevolella* species are involved in the pathogenesis of arthritis. While the intestinal flora was involved in the pathogenesis, occurrence and development of arthritis.

We also found enteritis in CIA model. At the acute stage of inflammation, ileitis is more serious than colitis, but in the recovery stage, intestinal inflammation basically disappeared, while colitis still exists. We also found that small intestinal inflammation in acute phase, and rapid regression; colitis is relatively mild, but lasting for a long time.

The pathological results confirmed that CII could cause intestinal inflammation in mice at the same time. The small intestinal inflammation lasted for a short time but more severe. Obvious inflammatory cell aggregation and tissue necrosis were observed in the mucosa and submucosa of small intestine. However, the colitis was relatively light, but the duration was longer than ileitis. The pathological results showed that there was local colonic inflammation two months after the second immunization. There were no obvious diarrhea and other symptoms in the whole experimental process. Whether the inflammation caused by collagen has an impact on intestinal flora and the changes of probiotics need further study.

The pathogenesis of RA is characterized by activation of macrophages by autoreactive T cells, resulting in the release of a series of pro-inflammatory cytokines. Saleem N etc [13] found that CII has has an important relationship with RA by T cell reactivity. Maybe the T cells and cytokines have an important role between microtiota and arthritis.

In conclusion, CII can induce arthritis and digestive system inflammation, such as enteritis, and may also induce respiratory, reproductive, circulatory and other system inflammation, we can name it collagen induced inflammations (CIIs), which is worthy of further study.

## COMPETING INTERESTS STATEMENT

The authors declare that they have no known competing financial interests or personal relationships that could have appeared to influence the work in this paper.

## REFERENCE

1. Liu S, Kiyoi T, Takemasa E, Maeyama K (2015) Systemic lentivirus-mediated delivery of short hairpin RNA targeting calcium release-activated calcium channel as gene therapy for collagen induced arthritis. J Immunol 194:76–83.

2. McInnes IB, Schett G (2011) The pathogenesis of rheumatoid arthritis. N Engl J Med 365:2205–2219.

3. Bessis N, Decker P, Assier E, Semerano L, Boissier MC (2017) Arthritis models: usefulness and interpretation. Semin Immunopathol 39:469–486.

4. Trentham DE (1982) Collagen arthritis as a relevant model for rheumatoid arthritis. Arthritis Rheum 25:911–916.

5. Kannan K, Ortmann RA, Kimpel D. Animal models of rheumatoid arthritis and their relevance to human disease. Pathophysiology, 2005;12:167–181.

6. Moudgil KD, Kim P, Brahn E. Advances in rheumatoid arthritis animal models. Curr Rheumatol Rep 2011;13:456.

7. Trentham DE, Townes AS, Kang AH. Autoimmunity to type II collagen an experimental model of arthritis. J Exp Med 1977;146:857–868.

8. Kannan K, Ortmann RA, Kimpel D. Animal models of rheumatoid arthritis and their relevance to human disease. Pathophysiology 2005;12:167–181.

9. D.D. Brand, K.A. Latham, E.F. Rosloniec, Collagen-induced arthritis, Nat Protoc. 2(5) (2007) 1269–1275.

10. K.S. Nandakumar, L. Svensson, R. Holmdahl, Collagen type II-specific monoclonal antibody-induced arthritis in mice: description of the disease and the influence of age, sex, and genes, Am. J. Pathol. 163 (2003) 1827–1837.

11. L.A. Joosten, E. Lubberts, P. Durez, M.M. Helsen, M.J. Jacobs, M. Goldman, W.B. van den Berg, Role of interleukin-4 and interleukin-10 in murine collagen-induced arthritis. Protective effect of interleukin-4 and interleukin-10 treatment on cartilage destruction, Arthritis Rheum. 40 (2) (1997) 249–260.

12. Maeda Y, Takeda K. Role of Gut Microbiota in Rheumatoid Arthritis. J Clin Med. 2017 Jun 9;6(6):60.

13. T cell reactivity to Collagen II as a possible prognostic marker in patients with rheumatoid arthritis. J Pak Med Assoc. 2018 Aug;68(8):1222–1227.

